# A Novel Bispecific Antibody CVL006 Superior to AK112 for Dual Targeting of PD-L1 and VEGF in Cancer Therapy

**DOI:** 10.1101/2024.12.05.627108

**Authors:** Chunyan Wang, Hao Huang, Zeng Song, Zhongyuan Li, Jinwen Huang, Liang Cao, Ziai Wu, Junfang Pan, Xiaokun Shen

## Abstract

Both preclinical and clinical studies have shown that combining anti-VEGF/VEGFR drugs with immune checkpoint inhibitors (ICIs) significantly enhances anticancer efficacy. Currently, PDL-1/ VEGF bispecific antibodies demonstrate superior antitumor activity compared to monotherapy or even the combination of PD-L1 inhibitors with anti-VEGF antibodies. This enhanced efficacy results from the simultaneous blockade of the PD-1/PD-L1 pathway and the inhibition of VEGF-driven angiogenesis. In this study, we developed a novel bispecific antibody, CVL006, by fusing an anti-PDL1 VHH domain with a humanized IgG1 anti-VEGF monoclonal antibody while retaining antibody-dependent cellular cytotoxicity (ADCC) functionality. CVL006 demonstrated high affinity and specificity for both human PD-L1 and VEGF, effectively blocking both the VEGF/VEGFR signaling pathway and the PD-L1/PD-1 axis. This dual blockade not only suppressed VEGF-induced angiogenesis but also reactivated T cells, increasing the secretion of cytokines essential for immune response. In vivo studies further indicated that CVL006 achieved superior antitumor efficacy compared to the PD-L1 inhibitor atezolizumab in mouse models, with greater tumor growth inhibition and reduced angiogenesis. To compare with approved bispecific antibody PD-1/VEGF AK112 (ivonescimab), CVL006 showed superior antitumor efficacy in vivo. These findings underscore the therapeutic potential of CVL006, which integrates immune checkpoint inhibition with disruption of tumor vascularization, offering a robust and comprehensive anticancer strategy.

## Background

Angiogenesis, primarily regulated by the VEGF/VEGFR signaling pathway, plays a critical role in both physiological and pathological processes (Bouis et al., 2006; Eelen et al., 2020). Bevacizumab, the first FDA-approved anti-VEGF monoclonal antibody, has been widely used to treat various cancers, including non-small cell lung cancer (NSCLC), metastatic breast cancer, and colorectal cancer (Garcia et al., 2020). Concurrently, antibodies targeting immune evasion, particularly those against PD-1 or PD-L1, have shown significant success in reactivating T cells to combat cancer (Ray et al., 2015). Clinical studies have demonstrated that anti-PD-1/PD-L1 antibodies, such as atezolizumab (Tecentriq), offer strong and durable anti-cancer effects in a range of malignancies, including lung cancer, renal cell carcinoma, melanoma, hepatocellular carcinoma, and lymphoma (Rizvi et al., 2015; Tsai and Daud, 2015; Beckermann et al., 2017; Li et al., 2022; Cao et al., 2023). Atezolizumab was the first PD-L1-targeting immunotherapy approved by the FDA for the treatment of urothelial carcinoma in 2016 (Ning et al., 2017). However, due to the complexity of cancer biology and interactions within the tumor microenvironment, single-target therapies often exhibit limited efficacy, with response rates to PD-1/PD-L1 antibodies ranging from 10% to 40% (Robert et al., 2015; Darvin et al., 2018; Jiang et al., 2020). To address these limitations, combination therapies and bispecific antibodies (BsAbs) have emerged as promising strategies to enhance therapeutic outcomes.

Recent studies have demonstrated that VEGF/VEGFR signaling impacts immune cells by promoting Treg cell proliferation, inhibiting dendritic cell maturation, and inducing PD-L1 expression on dendritic cells (Motz and Coukos, 2011; Khan and Kerbel, 2018). Additionally, VEGF suppresses CD8^+^T cell proliferation and upregulates inhibitory checkpoints, including PD-1, TIM-3, LAG-3, and CTLA-4, contributing to T cell exhaustion (Voron et al., 2015). Tumor-associated immune cells also facilitate tumor angiogenesis (Stockmann et al., 2014). Blocking VEGF/VEGFR signaling has been shown to enhance the antitumor efficacy of anti-PD-1/PD-L1 antibodies in preclinical models of colorectal, pancreatic, breast, and lung cancers (Yasuda et al., 2013; Voron et al., 2015; Allen et al., 2017; Meder et al., 2018). Clinical studies support the combination of atezolizumab and bevacizumab in improving outcomes for NSCLC patients, though this approach is associated with increased toxicity risks, such as fever, neutropenia, nephritis, and immune thrombocytopenic purpura, as observed in an analysis of 15,872 NSCLC patients (Reck et al., 2019). This underscores the need for novel strategies to enhance PD-(L)1 and VEGF blockade while minimizing toxicity.

AK112 is a first-in-class, humanized tetravalent bispecific antibody that simultaneously targets PD-1 and VEGF-induced tumor angiogenesis. AK112 offers a more comprehensive cancer treatment strategy compared to monotherapies or combination therapies targeting these pathways individually. Approved in May 2024 in China, AK112 combination with pemetrexed and carboplatin, is used to treat EGFR-mutated, locally advanced, or metastatic non-squamous NSCLC after progression on tyrosine kinase inhibitor (TKI) therapy. This approval followed the HARMONi-2 clinical trial, which demonstrated significant improvements in progression-free survival (PFS) and overall response rates compared to treatments like pembrolizumab. With a favorable safety profile and proven efficacy across multiple cancer types, AK112 is positioned as a next-generation therapeutic option. Ongoing trials, such as HARMONi-3, are further exploring its effectiveness in other cancers, reinforcing its potential as a versatile and potent treatment. As more data emerges, AK112 is expected to play a pivotal role in advancing immunotherapy.

Given the coexpression of VEGF and PD-(L)1 in the tumor microenvironment, dual PD-L1 and VEGF blockade promotes vascular normalization and enhances anti-tumor immune responses. We designed a novel bispecific antibody CVL006 combined an anti-PD-L1 VHH domain with a humanized IgG1 anti-VEGF monoclonal antibody. Through the analysis of its structure, in vitro affinity, functionality, and in vivo anti-tumor activity relative to AK112, we aimed to emphasize the unique features of CVL006. Preclinical studies indicated that CVL006 holds significant clinical promise. Its innovative design further highlights the growing role of bispecific antibodies in cancer therapy, opening new avenues for simultaneously targeting tumor vascular and immune evasion. As CVL006 progresses through clinical development, its ability to enhance cancer treatment outcomes will be rigorously evaluated, positioning it as a potentially valuable addition to the expanding arsenal of immunotherapy strategies.

## Experimental Methods

### Materials and reagents

The antibodies used in this study included anti-human PD-L1 (atezolizumab) from Genewiz (Suzhou, China) and anti-human VEGF (bevacizumab) from Sangon Biotech (Shanghai, China). CVL006 and human IgG1 isotype control antibody were synthesized by Wuxi Biologics (Shanghai, China). Recombinant proteins, including both human and mouse PD-L1, PD-1, VEGF, as well as PD-L1-expressing CHO-K1 cell lines, were prepared by Wuxi Biologics. Human umbilical vein endothelial cells (HUVECs) were purchased from ScienCell (Catalog No. #8000), and human IFN-γ antibodies were obtained from ThermoFisher (Rockford, IL, USA).

### Kinetics analysis of VEGF and PD-L1 binding

ELISA for VEGF binding: ELISA plates (Nunc MaxiSorp) were coated with 0.25 μg/mL human VEGF (Sino Biological, WBP325-hPro1) in carbonate-bicarbonate buffer (20 mM Na□CO□, 180 mM NaHCO□, pH 9.2) and incubated overnight at 4°C. The plates were blocked with 2% BSA for 1h at room temperature. Serial dilutions of CVL006 (0.001-10 μg/mL) were applied, incubated for 2h, and washed three times with PBS containing 0.5% Tween 20. Following this, 100 ng/mL Goat anti-human IgG Fc-HRP was added and incubated for 1h. Tetramethylbenzidine (TMB) substrate was used for staining, and reactions were stopped after 8 minutes by adding 2 M HCl. Absorbance was read at 450 nm using a SpectraMax® M5e reader, and EC_50_ values were determined with GraphPad Prism.

Flow cytometry (FACS) for PD-L1 Binding: CHO-K1 cells expressing human PD-L1 (WBP315.CHO-K1.hPro1.C11) were seeded in U-bottom 96-well plates at 1×10□ cells per well and incubated with serial dilutions of CVL006 (0.01-10 μg/mL) for 1h at 4°C. Following incubation, cells were washed with PBS or 1% BSA, and R-PE Goat anti-human IgG was added. After an additional 1h incubation in the dark, flow cytometry was performed to measure mean fluorescence intensity (MFI). EC_50_ values were calculated using FlowJo and GraphPad Prism.

### Surface plasmon resonance (SPR) for binding kinetics

The binding kinetics of CVL006 to VEGF and PD-L1 were measured using a Biacore T200 SPR system with a CM5 Sensor Chip. Anti-human Fc IgG was immobilized on the sensor surface, and serial dilutions of PD-L1 or VEGF antigens (W315-hPro1.ECD.his, WBP325-hPro1.his) were added into both association and dissociation phases at 25°C. The chip surface was regenerated between measurements using 10 mM glycine (pH 1.5). Binding kinetic rate constants (K_a_, K_d_) and equilibrium dissociation constants (KD) were calculated with Biacore T200 Evaluation software.

### Dual binding of VEGF and PD-L1

To assess bispecific binding, ELISA plates were coated with either VEGF or PD-L1 antigen (1μg/mL) in carbonate-bicarbonate buffer and incubated overnight at 4°C. Serial dilutions of CVL006 were applied, incubated for 1h at room temperature, followed by washing with PBS or 0.5% Tween 20. Biotinylated secondary antigen (VEGF-biotin or PD-L1-biotin) at μ0.5 g/mL was added for another 1h, followed by HRP-labeled streptavidin for detection. After TMB substrate development and reaction stoppage with 2 M HCl, absorbance was measured at 450 nm, and IC_50_ values were calculated using GraphPad Prism.

### HUVECs proliferation inhibition assay

HUVECs were seeded at 5000 cells/well in ECM containing 5% FBS and 1% ECGS. Cells were treated with 50 ng/mL VEGF and serial dilutions of CVL006 (0.001-10 μg/mL) and incubated at 37°C for 72h. Cell viability was measured with CellTiter-Glo, and IC_50_ values were calculated by plotting luminescence values against antibody concentration.

### Reporter gene assays

CHOK1-OKT3-PD-L1 cells were plated in 96-well plates at 20,000 cells/well and treated with serial dilutions of CVL006 and PD-1 expressing Jurkat cells containing an NFAT-RE-Luc2p reporter gene. After 4h, One-Glo luciferase assay reagent was added to measure luminescence.

### Cytotoxicity assays

HCC4006 cells were seeded at 8000 cells/well in 96-well plates and co-incubated with CVL006 and Jurkat-NFAT-Luciferase-CD16 effector cells at an 8:1 ratio for 6h at 37°C. Luminescence was measured after adding luciferase detection reagent. HCC4006 cells were washed, resuspended in Opti MEM, and incubated with antibody dilutions and 20% human AB serum for 3h at 37°C. After incubation, CellTiter-Glo reagent was added to measure cell viability and calculate IC_50_ values.

### In vivo tumor efficacy models

All animals were housed in a specific pathogen-free environment at WuXi Biologics Co., Ltd., with all experimental procedures approved by the company’s Animal Use and Care Committee.

In the hPBMC xenograft model, A375 tumor cells were washed with PBS, resuspended in a 1:1 mixture of PBS and Matrigel®, and adjusted to a concentration of 2×10□ cells/mL. For tumor establishment, each female NCG mouse (Nanjing GemPharmatech Co., Ltd.) was injected 2×10□ cells into the right flank. To support human tumor cell growth, 4×10□ human peripheral blood mononuclear cells (hPBMCs) from a healthy donor were injected into each mouse four days prior to tumor inoculation. Once tumors developed, mice were randomized into treatment groups (n=5-8 per group) based on initial tumor volume (TV), with average TVs of 80 mm^3^ for A375 tumors. Treatment groups included vehicle (PBS), CVL006 (6 mg/kg), atezolizumab (5 mg/kg), paclitaxel (10 mg/kg), and CVL006 combined with paclitaxel (10 mg/kg, respectively). Treatments were administered intraperitoneally three times a week, except for paclitaxel, which was given as a weekly intravenous injection. Tumor dimensions were recorded using a digital caliper, and tumor volume was calculated as TV = 0.5×a×b^2^, where a and b represent the longest and shortest tumor diameters, respectively. Tumor growth inhibition (TGI) was determined using the formula TGI = (1-T/C) × 100%, where T and C represent the average relative tumor volumes (RTV) of treated and control groups, respectively, with RTV calculated as the post-treatment tumor volume relative to baseline.

Given that CVL006 could not bind to murine VEGF, we substituted the VH and VL regions of bevacizumab with a newly designed CVL006-Surrogate antibody capable of targeting murine VEGF. For the colon adenocarcinoma (Colon 26) syngeneic tumor model, 5×10□ Colon 26 tumor cells were subcutaneously inoculated into the right flank of BALB/c mice. After 6 days, mice were randomly divided into 4 groups, and received intraperitoneal injections three times weekly of vehicle, atezolizumab+αVEGF (0.5+0.5 mg/kg), and CVL006-Surrogate (1.1 and 11.3 mg/kg).

For the standard xenograft model, female Balb/c nude mice were injected subcutaneously in the right flank with 5×10□ cells from different tumor cell lines, including OVCAR-3 and NCI-H1048. Once tumors reached an average volume of 100-200 mm^3^, mice were randomly assigned to treatment groups and received intraperitoneal injections three times weekly of vehicle, CVL006 (1.1 and 11.3 mg/kg), and AK112 (10 mg/kg; Akeso, Inc., Guangzhou, China).

### Statistical Analysis

Data were presented as mean ± SEM. The comparison between groups were performed using t-test or one-way ANOVA followed by uncorrected Fisher’s LSD. Statistical analyses were performed using GraphPad Prism 5. *P*<0.05 was considered to indicate statistical significance (^*^*P*<0.05, ^**^*P*<0.01, and ^***^*P*<0.001).

## Results

### Structure and affinity difference between CVL006 and AK112

Compared to AK112, CVL006 offers distinct advantages, primarily due to its refined structural design and retention of immune effector functions. Both antibodies are tetravalent bispecific based on bevacizumab; however, CVL006 uniquely incorporates an αPD-L1 VHH domain fused at the N-terminus of the light chain, enabling simultaneous targeting of PD-L1 and VEGF (Figure 1A-B). In contrast, prior studies have shown that AK112 employs an anti-PD-1 scFv at the C-terminus of the heavy chain, with Fc silencing mutations that significantly impair immune effector activities such as ADCC, complement-dependent cytotoxicity (CDC), antibody-dependent cellular phagocytosis (ADCP), and cytokines release (Zhong et al., 2023). CVL006, by maintaining an intact Fc region, preserves FcγR/IIIa binding, thereby facilitating stronger ADCC activity and promoting enhanced immune-mediated tumor cell elimination. This suggests that CVL006 may offer superior efficacy in cancers where robust immune activation is crucial, while AK112 focuses on reducing immune-related toxicity by limiting effector function.

**Figure 1.**
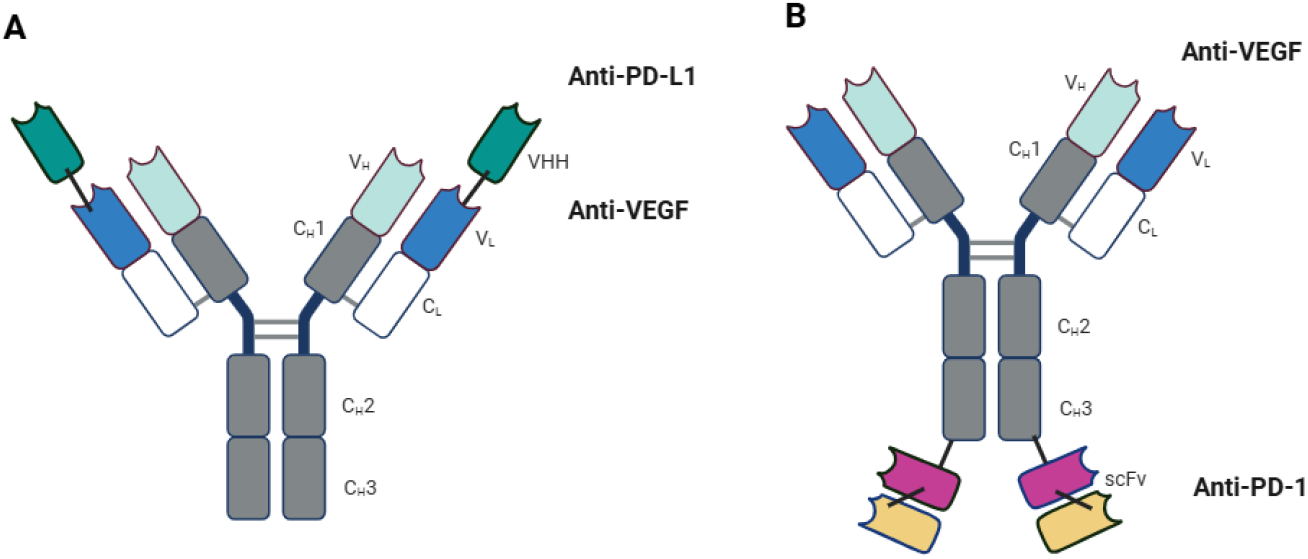
The structure diagram of CVL006 and AK112. (A) CVL006. (B) AK112.

The binding affinities of CVL006 to VEGF and PD-L1 were evaluated through ELISA and FACS. CVL006 exhibited comparable VEGF binding to bevacizumab, with an EC_50_ of 0.015 nM (Supplementary Table 1, Supplementary Figure 1). Similarly, in FACS assays, CVL006 demonstrated binding to PD-L1, comparable to the parent αPD-L1 VHH and atezolizumab, with an EC_50_ of 0.128 nM (Supplementary Table 1, Supplementary Figure 2).

SPR technology was employed to measure the affinity constant, as well as the on-rate and off-rate constants. Corrected experimental results showed KD values of 0.155 nM for PD-L1 and 0.015 nM for VEGF, indicating that CVL006 exhibited binding affinities comparable to its parent antibodies (Supplementary Table 1, Supplementary Figure 3A and 3B). As showed in Supplementary Table 1, CVL006 outperformed AK112 in VEGF binding activity, with more than 2X and 21X higher potency in ELISA and SPR assays, respectively (Zhong et al., 2022). Additionally, in FACS and SPR assays, CVL006 demonstrated superior binding affinity to AK112, by more than 27X and 1.5X, respectively. These data strongly suggest that CVL006 may possess superior therapeutic potential relative to AK112, and direct head-to-head comparative studies need to explore in future.

### VEGF efficiently enhanced the binding of CVL006 to PD-L1

With published data of AK112, the binding affinity of AK112 to PD-1 increased more than 10-fold when VEGF was present, enhancing its potency in blocking PD-1/PD-L1 signaling and increasing T-cell activation in vitro (Zhong et al., 2022). This finding highlighted the role of VEGF in amplifying the therapeutic potential of bispecific antibodies. To explore whether VEGF could exert a similar enhancement on CVL006’s interaction with PD-L1, we conducted binding assays using ELISA in the presence of VEGF.

As shown in Figure 2, VEGF significantly boosted CVL006’s binding to PD-L1, reducing the EC_50_ from 0.623 nM to 0.048 nM—an approximate 13-fold improvement in binding efficiency. This dramatic increase in affinity suggests that CVL006, like AK112, benefits from the presence of VEGF, which facilitates more accurate and efficient targeting of PD-L1-positive tumor cells. This interaction likely increases the concentration of CVL006 within the tumor microenvironment, where VEGF is highly expressed, enhancing its tumor specificity and minimizing off-target effects. These data further support the hypothesis that CVL006’s dual targeting of VEGF and PD-L1 could lead to improved anti-tumor efficacy by leveraging VEGF’s presence to increase binding precision and functional activity within tumors.

**Figure 2.**
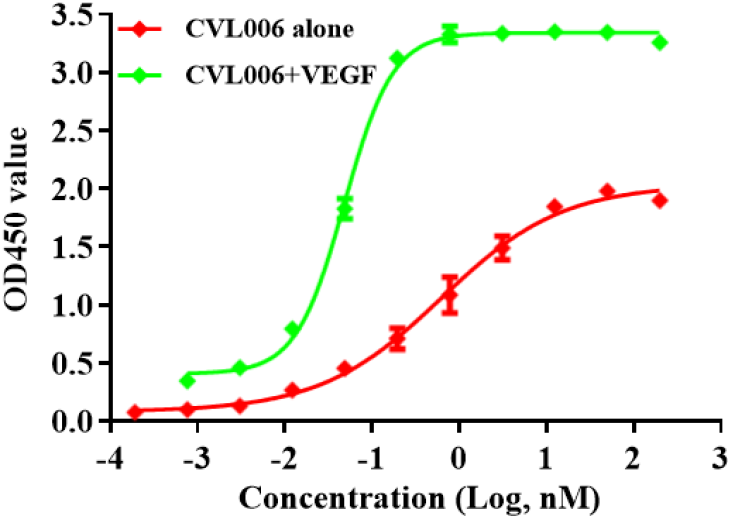
The effect of VEGF on binding affinity of CVL006 to PD-L1.

### Functional assay in VEGF-mediated HUVECs proliferation and PD-L1/PD-1 signal pathway

To compare the biological activities of CVL006 and AK112, we assessed their inhibitory effects on VEGF-induced proliferation of HUVECs (IC_50_ values are 5.3, 4.0, and 3.0 nM, respectively). Although results demonstrated that CVL006 is comparable with AK112 in suppressing HUVEC proliferation, as this cell growth assay may lack precision (Figure 3A). Additionally, to investigate their inhibitory effects on the PD-1/PD-L1 signaling pathway, we employed a luciferase reporter gene assay to measure NFAT activation. In this assay, CHOK1-OKT3-PD-L1 cells (PD-L1-overexpressed target cells) were co-cultured with NFAT-RE-Luc2p-integrated, full-length PD-1-expressed Jurkat cells (activated effector T cells). The data revealed that CVL006, AK112, and atezolizumab demonstrated similar potencies in blocking PD-1/PD-L1 signaling, with IC_50_ values of 6.9, 3.1, and 4.1 nM, respectively (Figure 3B).

**Figure 3.**
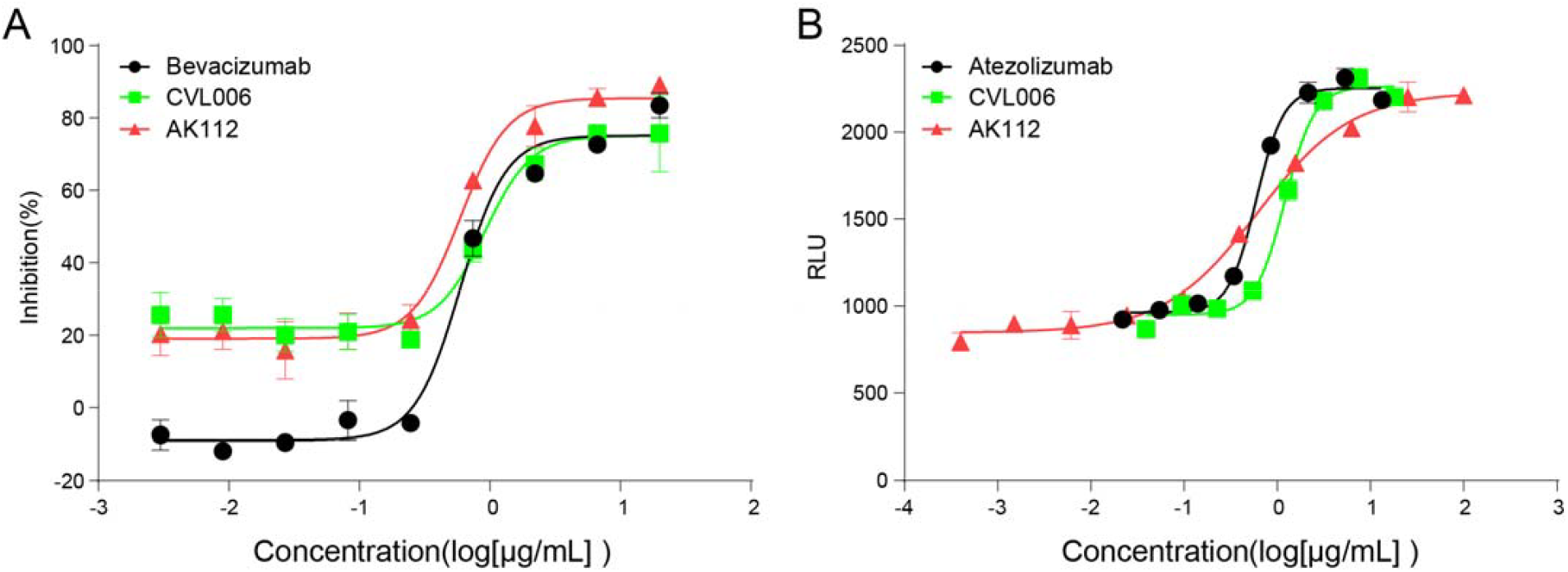
The biological activity of antibody to VEGF/VEGFR and PD-L1/PD-1 signal pathway. (A) The inhibition effect of CVL006, bevacizumab, and AK112 on VEGF-induced HUVEC proliferation assay. (B) Blockage effect of CVL006, atezolizumab and AK112 on the PD-L1/PD-1 pathway by the luciferase reporter gene system.

These findings indicate that CVL006 exhibits comparable biological activity to AK112 in both VEGF inhibition and PD-1/PD-L1 pathway blockade. This supports its potential as a therapeutic agent capable of targeting both tumor vascular and immune evasion mechanisms.

### CVL006 showed ADCC effect

The IgG1 isotype is well known for its potent effector functions. Using the human lung adenocarcinoma cell line HCC4006 as the target and stably CD16-expressed Jurkat NFAT luciferase reporter cells as effector cells, the ADCC activities of bevacizumab and CVL006 were evaluated through luciferase signaling assays. As shown in Figures 4A-B, bevacizumab exhibited undetectable ADCC activity, consistent with the hypothesis that bevacizumab neutralizes VEGF in solution rather than on the cell surface. In contrast, CVL006 displayed significant ADCC activity against HCC4006 cells, with an EC_50_ of 0.2436 μg/mL, demonstrating its potential to engage effector cells for immune-mediated tumor cell killing.

**Figure 4.**
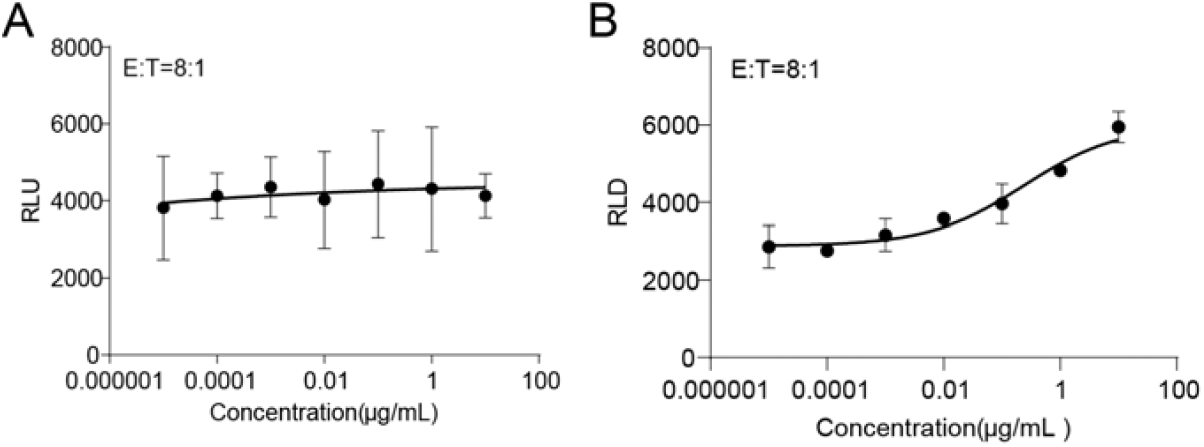
The ADCC effect of CVL006 on HCC4006 cells. (A) The ADCC effect of Bevacizumab. (B) The ADCC effect of CVL006. Bevacizumab was used as negative controls.

In comparison, AK112 was specifically engineered to eliminate ADCC through Fc silencing mutations, thereby avoiding potential damage to PD-1-positive lymphocytes. This design is intended to minimize the risk of off-target effects, particularly in immune cells, while maintaining its therapeutic efficacy. These findings suggest that while CVL006 retains the ability to engage immune effector cells and promote ADCC, AK112 prioritizes safety by eliminating such activity to protect immune cells, thus offering distinct advantages depending on the therapeutic context.

### CVL006 significantly inhibited tumor growth

In the colon adenocarcinoma (Colon 26) syngeneic tumor model, TGI rates were assessed on day 15 post-treatment. The vehicle, atezolizumab+αVEGF, and CVL006-Surrogate (administered at 1.1 and 11.3 mg/kg) achieved TGI rates of 43.18%, 48.52% and 67.69%, respectively, all demonstrating significant tumor growth suppression compared to the control group. CVL006-Surrogate exhibited superior anti-tumor efficacy compared to atezolizumab, with enhanced activity observed at the higher dose. No significant changes or signs of toxicity were observed throughout the study (Figure 5A).

**Figure 5.**
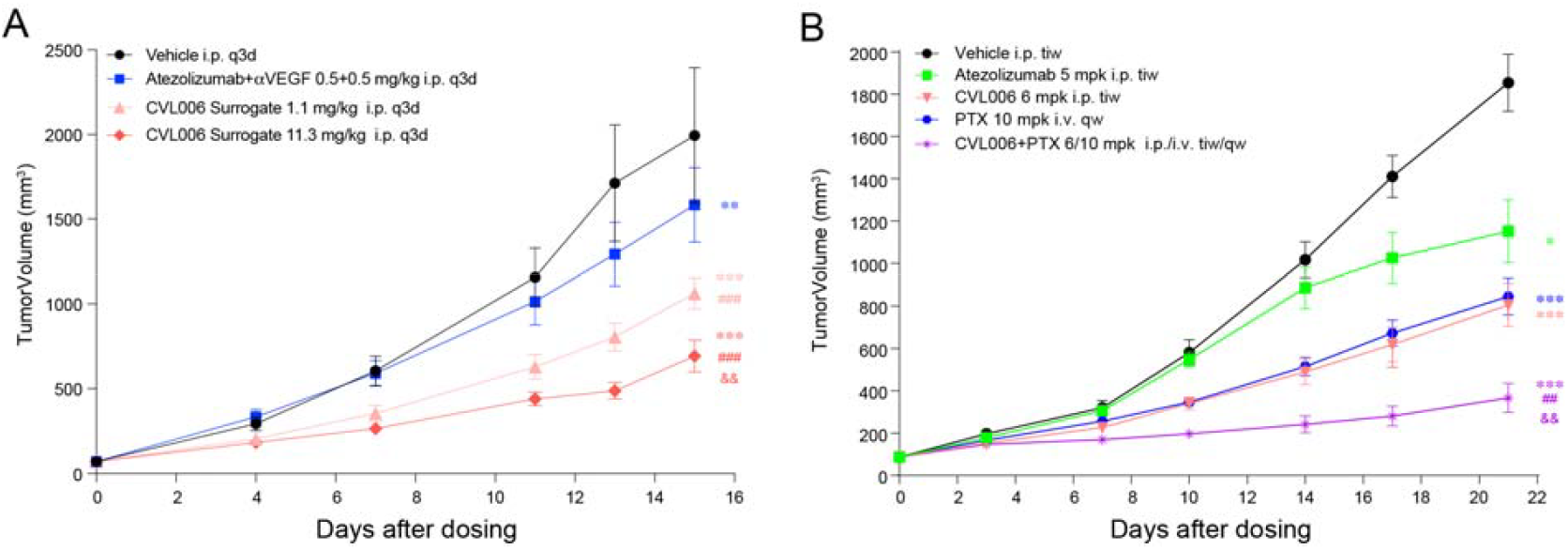
The in vivo efficacy of CVL006 was evaluated in Colon 26 syngeneic tumor model and hPBMC+A375 xenograft model. (A) Colon 26 syngeneic tumor model. (B) hPBMC+A375 xenograft model. Data expressed as mean ± SEM. ^*^*P*<0.05 ^**^*P*<0.01, ^***^*P*<0.001 vs vehicle; ^##^*P*<0.01, ^###^*P*<0.001, vs Atrzolizumab+αVEGF in (A), vs CVL006-6 mg/kg in (B); ^&&^*P*<0.01, vs CVL006 Surrogate 1.1mg/kg in (A), PTX-10 mg/kg in (B).

In the hPBMC+A375 xenograft model, treatment with atezolizumab, bevacizumab, CVL006, paclitaxel, and the combination group significantly inhibited tumor growth, with TGIs ranging from 41.48% to 85.40% (*P*<0.001). CVL006 displayed a dose-dependent anti-tumor effect and showed greater potency compared to atezolizumab at equivalent doses, with a better safety profile. The combination of CVL006 with paclitaxel demonstrated a synergistic effect, further enhancing anti-tumor efficacy. No toxicity-related weight loss or side effects were observed (Figure 5B).

In additional xenograft models, including ovarian cancer (OVCAR-3) and lung cancer (NCI-H1048), CVL006 demonstrated robust anti-tumor activity comparable to AK112 in ovarian cancer. Strikingly, in the NCI-H1048 lung cancer xenograft model, CVL006 exhibited significantly superior efficacy compared to AK112, with tumor volume reductions indicating enhanced therapeutic potential (*P*<0.01) (Figure 6A and 6B). These experiments were conducted in immunodeficient mice, ensuring that the observed anti-tumor effects were predominantly mediated by VEGF neutralization. This suggests that CVL006’s superior performance in the NCI-H1048 model could be attributed to its enhanced VEGF blockade efficiency, highlighting its promise as a potent therapeutic candidate in VEGF-driven tumors.

**Figure 6.**
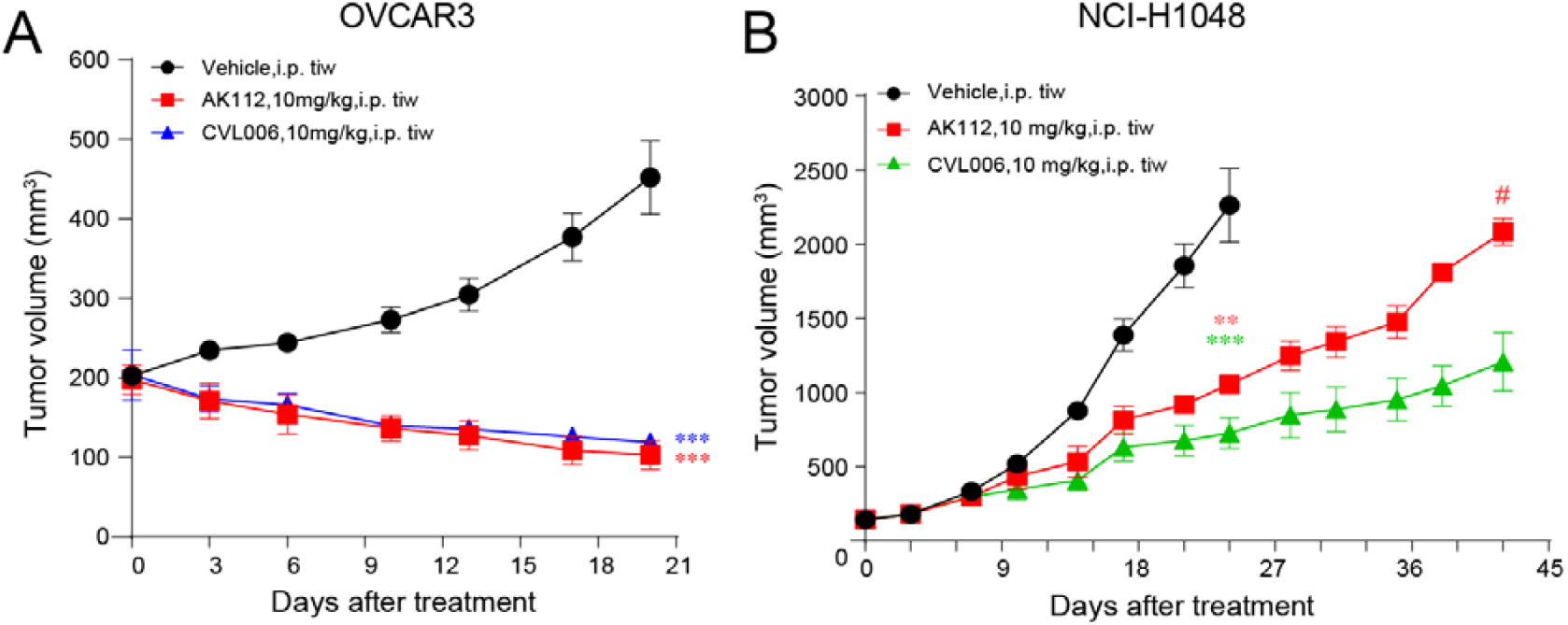
The in vivo efficacy of CVL006 was evaluated in xenograft models. (A) OVCAR-3, (B) NCI-H1048. All antibodies were administered intraperitoneally (i.p.) thrice a week (TIW). Data expressed as mean ± SEM. ^*^*P*<0.05 ^**^*P* 0.01<^***^*P*<0.001 vs vehicle.

Overall, CVL006 demonstrated strong potential as a therapeutic agent, with superior or comparable efficacy to AK112 in xenograft models, while maintaining a favorable safety profile. These results suggest further exploration of its mechanisms and potential advantages in immune-competent models or combination therapies.

## Discussion

The VEGF/VEGFR signaling pathway is commonly upregulated in tumors, promoting cancer cell proliferation and angiogenesis (Liu et al., 2023). VEGF contributes to an immunosuppressive tumor microenvironment by increasing regulatory T cells (Tregs), suppressing CD8^+^T cell function, and upregulating immune checkpoints such as PD-1, TIM-3, LAG-3, and CTLA-4 (Motz and Coukos, 2011; Voron et al., 2015; Khan and Kerbel, 2018). While immune checkpoint inhibitors (ICIs), such as anti-PD-1/PD-L1 therapies, have demonstrated clinical success with objective response rates (ORRs) ranging from 10-40% in solid tumors, tumor immune escape mechanisms often limit their efficacy (Darvin et al., 2018). The combination of anti-VEGF and ICIs has emerged as a promising strategy to enhance therapeutic outcomes and overcome resistance. Preclinical studies have shown that VEGF inhibitors, beyond their antiangiogenic effects, can synergize with ICIs by normalizing tumor vasculature, increasing T-cell infiltration, and reducing the activity of immunosuppressive cells like Tregs and myeloid-derived suppressor cells (Gabrilovich et al., 1998; Hegde et al., 2018; Hack et al., 2020).

Clinically, this approach has been validated in trials such as IMpower150, where the combination of atezolizumab, bevacizumab, and chemotherapy significantly improved survival in metastatic non-squamous NSCLC, and IMbrave150, where combining atezolizumab and bevacizumab improved overall survival (OS) and progression-free survival (PFS) in unresectable hepatocellular carcinoma (Socinski et al., 2018; Liu et al., 2019; Finn et al., 2020; Socinski et al., 2021). These findings highlight the synergistic potential of combining anti-VEGF and anti-PD-1/PD-L1 therapies, leading to enhanced anti-tumor efficacy across various solid tumors.

This study presents a novel IgG-like bifunctional molecule, CVL006, designed to inhibit PD-L1-mediated immunosuppression while simultaneously blocking VEGF-induced tumor angiogenesis within the tumor microenvironment (TME). Comprehensive in vitro and in vivo characterizations were conducted to assess its dual functionality. At the molecular level, SPR affinity assays confirmed that CVL006 effectively retained high binding affinities for both PD-L1 and VEGF. Functionally, as a PD-L1 inhibitor, CVL006 demonstrated comparable efficacy to its parental anti-PD-L1 antibody and the therapeutic antibody atezolizumab, effectively restoring T-cell activity. Furthermore, CVL006 retained comparable functional potency in blocking VEGF-mediated cell proliferation when assessed against bevacizumab, indicating its potential to suppress angiogenesis and reduce tumor progression. These findings highlight CVL006 as a promising therapeutic agent with dual mechanisms of action, offering potential advantages in targeting both immune evasion and tumor vascularization within the TME, thereby addressing two critical aspects of tumor progression.

PD-L1 blockade therapy, as a cornerstone of immune checkpoint inhibition, has demonstrated remarkable efficacy across various malignancies, particularly in immunologically active tumors such as lung cancer (Forde et al., 2018), melanoma (VanderWalde et al., 2023), hepatocellular carcinoma (HCC) (Lu et al., 2022), ovarian cancer (Fukumoto et al., 2019), and colorectal cancer, significantly improving overall survival and progression-free survival (Brahmer et al., 2012). However, the suboptimal response rates of PD-L1 monotherapy in certain patient populations have spurred interest in combination strategies, with particular attention to the synergy between PD-L1 and VEGF inhibitors (Yi et al., 2022). This dual approach leverages complementary mechanisms by suppressing tumor immune evasion and inhibiting angiogenesis, thereby addressing resistance while enhancing immune infiltration into the tumor microenvironment. In the NCI-H1048 lung cancer model, CVL006 has demonstrated superior antitumor activity compared to AK112, attributed to its higher PD-L1 binding affinity, potent immune activation capability, and enhanced synergy in angiogenesis inhibition. CVL006 significantly delayed tumor growth, reduced angiogenesis, and favorably modulated the immune microenvironment, highlighting its potential as a next-generation PD-L1 inhibitor. These findings not only provide a strong foundation for the clinical development of CVL006 but also underscore the promise of PD-L1 and VEGF inhibitor combinations as a transformative strategy for improving treatment outcomes in cancer patients through precise and effective therapies.

Our study also demonstrated that the combination of CVL006 with the chemotherapy agent paclitaxel exhibited remarkable synergistic antitumor effects in the xenograft mouse model. This synergy likely arose from their complementary mechanisms: CVL006 effectively disrupted tumor immune evasion and inhibited angiogenesis, while paclitaxel directly targeted tumor cells by stabilizing microtubules and arresting cell division. The combined treatment achieved superior tumor growth suppression compared to monotherapy. Clinically, this combination therapy holds considerable promise, offering the potential to enhance efficacy at reduced doses, thereby minimizing chemotherapy-induced toxicity and improving patient tolerability. Moreover, this approach could represent a valuable therapeutic option for patients with resistance to monotherapy. These findings provide a solid foundation for the clinical development of CVL006 and paclitaxel combination therapy; however, further investigations are needed to optimize dosing strategies, elucidate underlying mechanisms, and validate safety and efficacy in clinical trials.

In summary, the rational structural design, high specificity in target affinity, and robust in vivo anti-tumor activity of CVL006 collectively highlight its strong potential to become a best-in-class therapeutic agent. These features position CVL006 as a promising candidate for targeting both immune evasion and tumor angiogenesis, addressing key mechanisms involved in tumor progression.

---

**Supplementary table 1.**
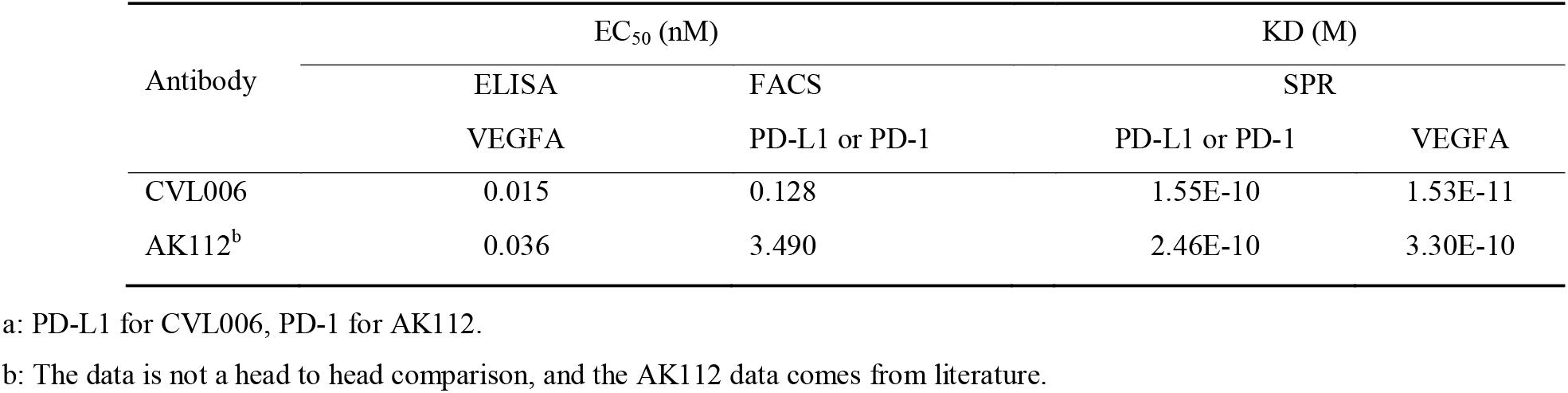
Antigen binding activity of CVL006 and AK112.

**Supplementary table 2.** Kinetic affinity results of antibodies.

| Analyte | Ligand | ka (1/Ms) | kd (1/s) | KD (M) |
| --- | --- | --- | --- | --- |
| PD-L1(W315-hPro1.ECD.his) | CVL006 | 2.14E+06 | 3.33E-04 | 1.55E-10 |
|  | Anti-PD-L1 VHH | 1.82E+06 | 2.91E-04 | 1.60E-10 |
|  | Atezolizumab | 1.53E+06 | 9.29E-05 | 6.08E-11 |
| VEGF(W325-hPro1.ECD.his) | CVL006 | 2.38E+06 | 3.64E-05 | 1.53E-11 |
|  | Bevacizumab | 3.94E+06 | 3.69E-05 | 9.36E-12 |

**SFigure 1.**
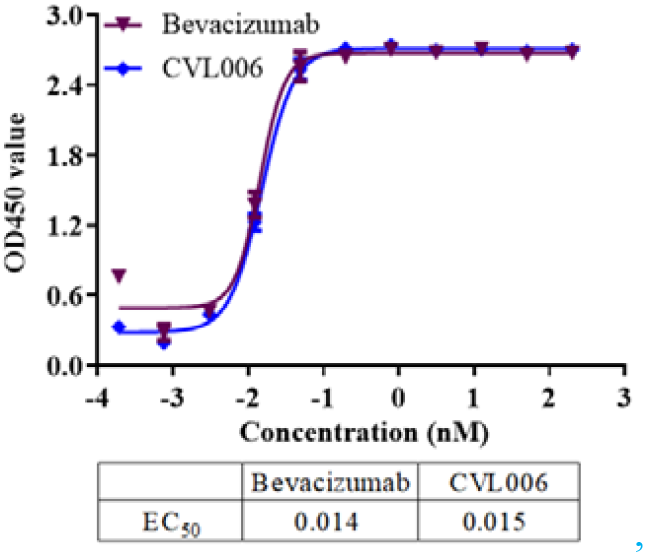
The ELISA binding activity of CVL006 and bevacizumab to human VEGF.

**SFigure 2.**
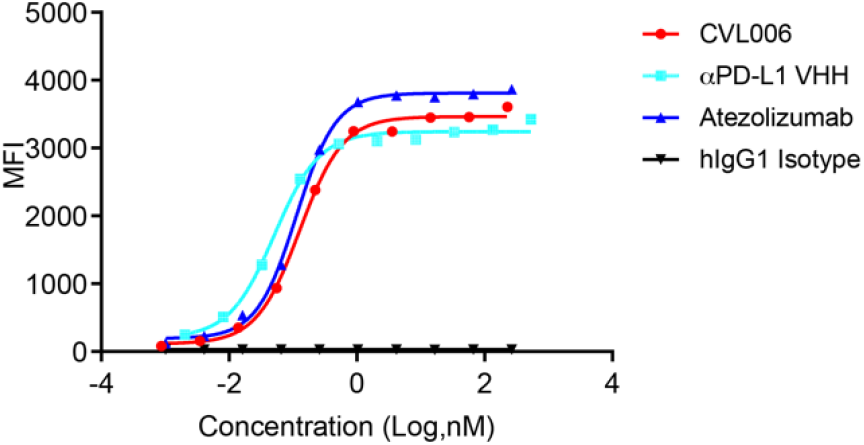
The FACS binding activity of CVL006, αPD-L1 VHH, and atezolizumab to PD-L1 expressing cell.

**SFigure 3.**
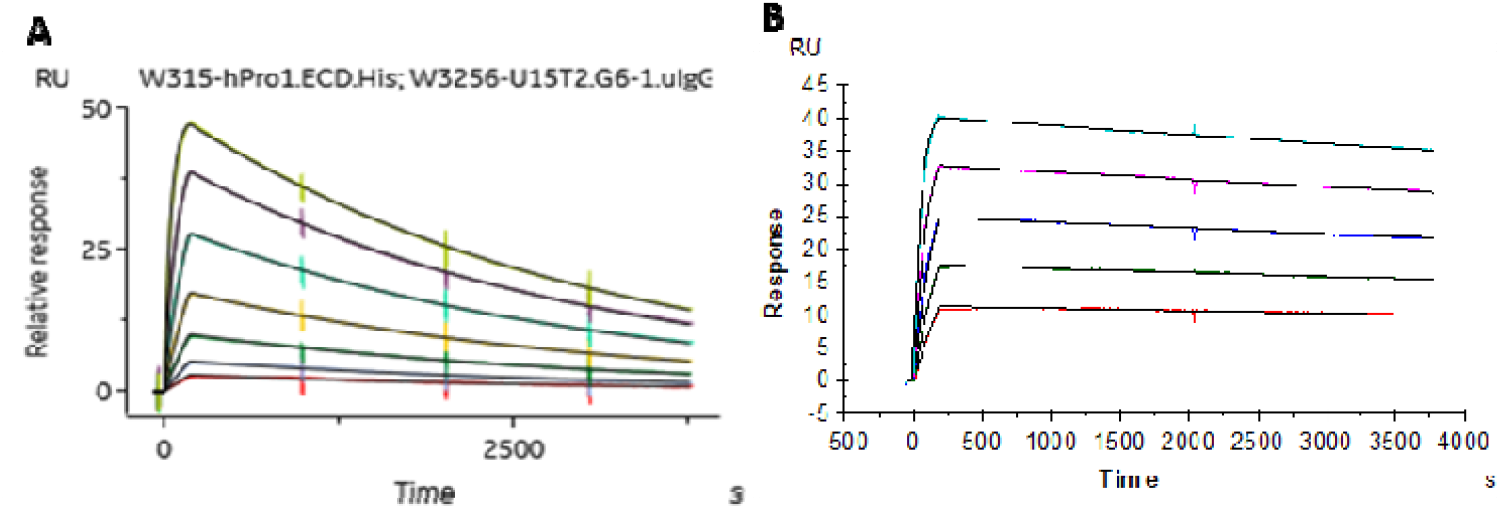
The kinetic binding profile of CVL006. (A) The SPR Sensorgram of CVL006 binding to human PD-L1. (B) The SPR Sensorgram of CVL006 binding to human VEGF.

